# The transcriptional landscape of human microglia reveals strong conservation of miRNAs and preservation of function across vertebrate species

**DOI:** 10.64898/2026.04.20.719771

**Authors:** Sarrabeth Stone, Alexander D. Walsh, Natalie Sol-Foulon, Lars Pennings, Elodie Martin, Liz Baretto Arce, Richard J. Leventer, Trevor J. Kilpatrick, Paul J. Lockhart, Bernard Zalc, Brendan R.E. Ansell, Michele D. Binder

## Abstract

The central role of microglia in CNS function in health and disease has resulted in large interest in targeting microglial as treatments for neurodegenerative disease; understanding the factors that regulate microglial gene expression will be crucial to this goal. microRNAs (miRNAs) are among the most abundant post transcriptional regulators of gene expression. miRNAs suggests miRNA were likely key to significant evolutionary events as regulators of gene expression. The miRNAome of microglia is critical to their correct functioning but the miRNA that define microglia identity and regulate key functions have not been fully defined. In this study we performed a detailed analysis of the microglial miRNAome to identify miRNA enriched in microglia that are conserved across species (human, mouse, and xenopus). We further characterised the expression of these conserved miRNAs during demyelination and remyelination and identified conserved function of a microglial–enriched miRNA across species. These findings reveal evolutionary conservation of specific miRNAs, suggesting an important role in establishing and maintaining microglial identity. They also highlight miRNAs that may be critical for microglial function in the central nervous system in both health and disease. Overall, this work advances our understanding of the factors that regulate microglial gene expression.

## Introduction

Microglia are a type of immune cell found exclusively within the CNS in vertebrates and some invertebrates^1^. Microglia arise outside the CNS and enter the CNS early in development^2,3^. Microglia are essential to the maintenance of brain homeostasis and the formation of neural connections via synaptic pruning^4^. Microglia have also been demonstrated to play a role in the development of oligodendrocytes, and ultimately in myelination^5–8^. Many of these functions are conserved across species suggesting the likelihood of a shared transcriptional programme^1^. Moreover, as the resident immune cell of the brain microglia have vital roles in the initiation and resolution of inflammatory responses and have been implicated in the pathogenesis of CNS diseases including Alzheimer’s, Parkinson’s, and multiple sclerosis (MS)^9,10^. The transcriptional programmes underlying the response of microglia to brain damage and disease are beginning to be elucidated^11^. However, relatively little attention has been paid to the role of small RNAs, such as microRNAs (miRNAs), in this process, despite their critical role in refining and regulating transcriptional programmes.

miRNAs are type of endogenous short non-coding RNA that negatively regulate gene expression via inhibiting translation or inducing degradation of mRNA^12^. First described as regulators of developmental timing in *C.elegans*^13,14^, miRNAs are widely conserved across vertebrate and invertebrate species^15,16^. More than 60% of mammalian protein coding genes contain motifs allowing them to be regulated by one or more miRNAs, and a single miRNA can target many mRNA transcripts^16,17^. The ability of a single miRNAs to interact with multiple genes allows miRNAs to act as ‘master regulators’ of the transcriptional response to intrinsic and extrinsic cues^18^.

Consistent with the role of miRNAs in defining cell identity, the miRNAome of microglia is distinct from other cell types^19–21^, and is vital for the normal function of microglia. Further, the miRNAome of microglia changes substantially across the lifespan of mice^21^. Microglial-specific deletion of Dicer in prenatal mice, which results in the loss of all microglial miRNAs and leads to inflammatory activation of microglia and DNA damage. However, this deletion does not result in the failure of microglia to migrate to the developing CNS^20^. Similarly, miRNAs in adult microglia are not required for homeostatic maintenance of the microglial population in mice but miRNA-deficient microglia are hyper responsive to inflammatory stimuli^20^. The transcriptome of human microglia has been shown to vary through ontogeny^22^, however, the extent to which the microglial miRNAome changes throughout ontogeny has not been previously studied in humans.

Activation of microglia to pro-inflammatory and anti-inflammatory-like phenotypes results in differential expression profiles of miRNA, suggesting that miRNAs are important regulators of microglial responses to CNS damage and disease^23^. Microglial miRNA deficiency has been demonstrated to impair hippocampal neuron function, increase pathology in a mouse model of tauopathy, and render microglia hyper-responsive to inflammatory stimuli^20,24^. Additionally, in the cuprizone model of demyelination microglial miRNA deficiency results in microglia hyperreactivity, increased demyelination, and impaired remyelination^25^. In people with MS, differential expression of miRNAs can be detected in peripheral immune cells and serum of ^26,27^. Studies of MS lesions also reveal dysregulation of miRNAs in the CNS^28–30^.

In this study, we performed a cross-species analysis of microglial miRNA expression comparing microglia from humans and two model organisms, and examined miRNA dynamics in the cuprizone mouse model of demyelination (Fig1). RNA sequencing of live human microglia purified from neurosurgical samples defined both the mRNA transcriptome and miRNAome. We identified a distinct human microglial miRNA profile and a conserved core miRNA signature shared by humans, mice and *Xenopus*. Analysis of the microglial miRNAome during cuprizone induced demyelination and remyelination identified microglial-enriched miRNAs that showed altered expression during demyelination and remyelination. Functional studies demonstrated that overexpression of one of these miRNAs, miR-511-3p, in primary mouse microglia or human iPSC-derived microglia-like cells resulted in decreased phagocytosis of myelin debris. Collectively, these finding demonstrate evolutionary conservation of microglial miRNA expression signatures and function and identify miRNAs which may be critical in regulating the microglial response to CNS damage.

## Results

### Human microglia express a unique pattern of miRNAs compared to other brain cells

Microglia express a unique profile of miRNA which play key roles in regulating microglial function^20,24^. However, studies of human microglial miRNA have been limited. Thus, we set out to understand the transcriptional miRNA landscape of human microglia and its relevance to cell function. Brain tissue was obtained from individuals (N=23) undergoing neurosurgery for lesions associated with intractable focal epilepsy (Supplementary Table 1). Normal appearing non-lesional tissue was separated into two cell populations; CD45^+ve^ microglia and CD45^-ve^ bulk brain cells using immunopanning, which enables the isolation of highly purified microglia^21^. We then performed miRNA and mRNA sequencing on these two populations (Fig.1).

**Figure 1.**
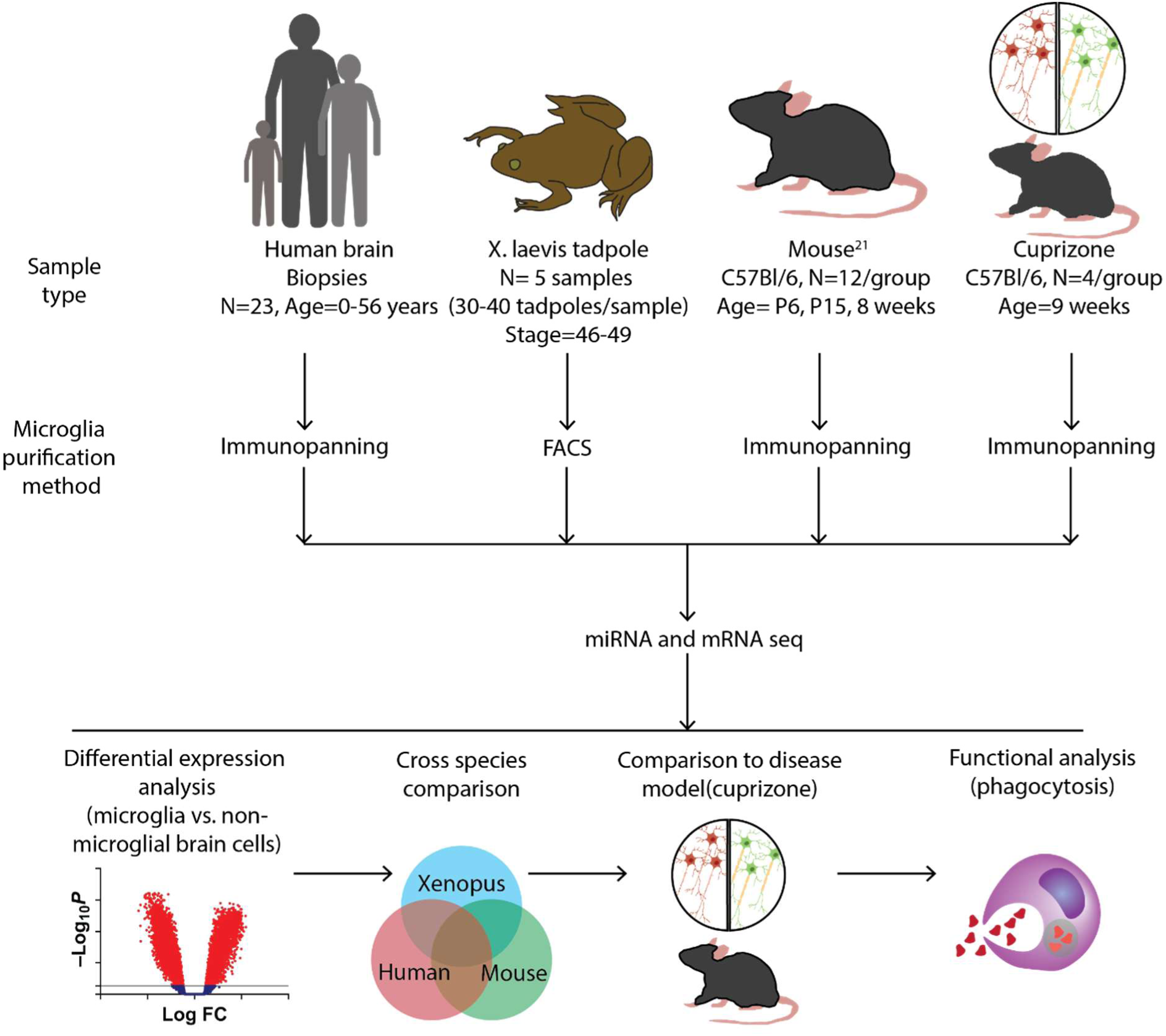
miRNA and mRNA experimental workflow.

Initial QC of the dataset resulted in the exclusion of some samples. One CD45^-ve^ bulk sample with fewer than 1 million reads was removed from the analysis. Purity of the CD45^+ve^ microglia population was then assessed using digital cytometry (CIBERSORTx) analysis comparing our RNASeq data with a single nucleus sequencing dataset generated from adult human brain samples^31^. Seven CD45^+ve^ microglia samples with <50% purity were excluded from further analysis (Supplementary Fig.1). Importantly, the CIBERSORTx analysis of the CD45^-ve^ bulk samples demonstrated comparable microglial depletion between all CD45^-ve^ bulk samples irrespective of CD45^+ve^ microglial population purity (2.3% high microglia purity vs 1.3% low microglial purity), therefore, all CD45^-ve^ bulk cell samples were retained in the analysis. The purity of the remaining CD45^+ve^ microglia samples was 79.6% ± 10.6 (mean ± SD); with non-microglial cells composed of neurons (6.7% ± 4.4), oligodendroglial lineage cells (9.5% ± 5.4), and astrocytes (4.3% ± 2.7). In contrast the CD45^-ve^ bulk cell samples were largely depleted of microglia (1.9% ± 2.4) and contained neurons (20% ± 13), oligodendroglial lineage cells (48.7% ± 21.1), and astrocytes (29.3% ± 13.3). Overall, 16 CD45^+ve^ microglial and 22 CD45^-ve^ bulk cell samples passed QC and proceeded for further analysis (Supplementary Table 1). Multidimensional scaling (MDS) analysis of miRNA and mRNA expression revealed a strong separation of the CD45^+ve^ microglial cells and CD45^-ve^ bulk brain cells indicating distinct expression signatures for these two cell populations (Fig.2A, B). Further MDS analysis showed that samples did not cluster based on disease, sex, or brain region (Supplementary Fig. 2).

**Figure 2.**
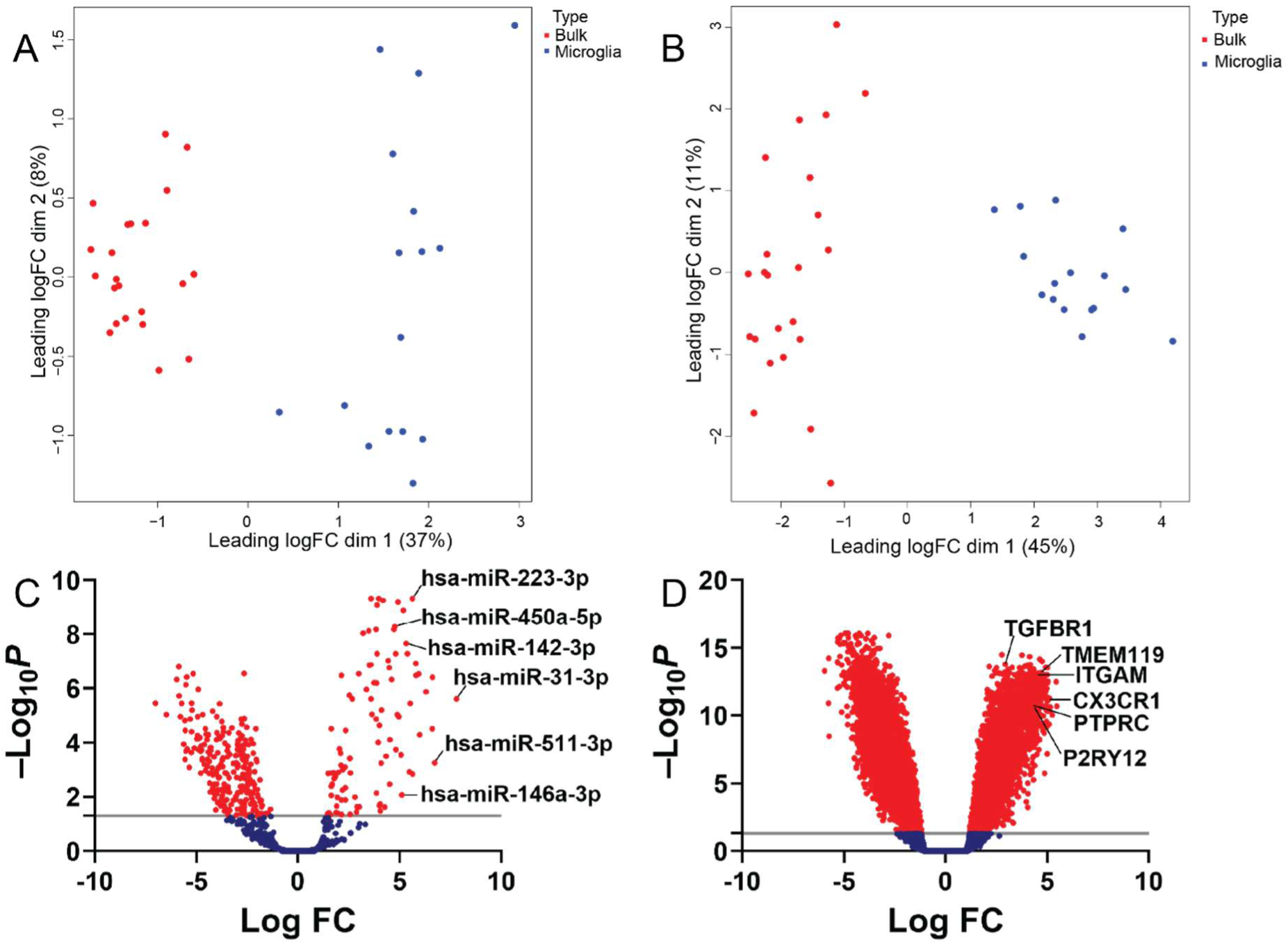
Human microglial have a unique miRNA transcriptional profile compared to non-microglial brain cells. **A.** MDA analysis of miRNA sequencing data from CD45^+ve^ and CD45^-ve^ cell types reveals strong clustering by cell type. **B.** MDA analysis of mRNA sequencing data from CD45^+ve^ and CD45^-ve^ cell types reveals strong clustering by cell type. **C.** Volcano plot of miRNA that were differentially expressed between pooled CD45^+ve^ microglia and CD45^-ve^ bulk brain cells. 97 of 838 miRNA were enriched in CD45^+ve^ microglia compared to CD45^-ve^ bulk brain cells (log2FC > 1, adj.P.Val <0.05). **D.** Volcano plot of mRNA that were differentially expressed between pooled CD45^+ve^ microglia and CD45^-ve^ bulk brain cells. 3254 of 23,378 mRNA were enriched in CD45^+ve^ microglia compared to CD45^-ve^ bulk brain cells (log2FC > 1, adj.P.Val <0.05). N= 16 CD45^+ve^, 22 CD45^-ve^.

Sequencing identified 838 miRNA transcripts, and 23,378 mRNA transcripts (Fig.2C, D). We compared the expression profiles of CD45^+ve^ microglia to those of CD45^-ve^ bulk brain cells to identify miRNA and mRNA transcripts enriched in microglia. We identified 97 miRNAs and 3254 mRNAs significantly enriched in microglia compared to the CD45^-ve^ bulk brain cells (log2FC > 1, adj.P.Val <0.05) (Fig.2C, D, Supplementary Tables 2 and 3). Notably, known microglial markers including *CX3CR1*, *TREM2*, *TMEM119*, hsa-miR-511-3p, hsa-miR-223-3p, and hsa-miR-142-3p were significantly enriched, further validating the effectiveness of the immunopanning protocol^5,21^. This rich dataset extends our knowledge of the miRNAome of human microglia.

### Xenopus microglial miRNA

Microglia promote myelination and remyelination in the CNS^5,7,8,32,33^. To identify genes and pathways that may contribute to these processes, we performed cross-species analysis using our human dataset, a mouse microglia dataset we previously generated^21^, and the laboratory model *Xenopus laevis.* To characterise the transcriptional landscape of *X. laevis* microglia, we used transgenic *X. laevis* tadpoles expressing mCherry under the control of the MPEG promotor, which is active in macrophage lineage cells, including microglia^34,35^. Brains were harvested from stage 46–49 tadpoles (N=5), which corresponds to myelination in the *Xenopus* brain^36^. mCherry^+ve^ microglia were separated from mCherry^-ve^ bulk brain cells by FACS (Supplementary Fig.3) and both the miRNA and mRNA from these populations were sequenced. MDS analysis of miRNA and mRNA expression revealed distinct clustering of mCherry^+ve^ microglia and mCherry^-ve^ bulk brain cells, indicating unique expression profiles (Fig.3A, B).

**Figure 3.**
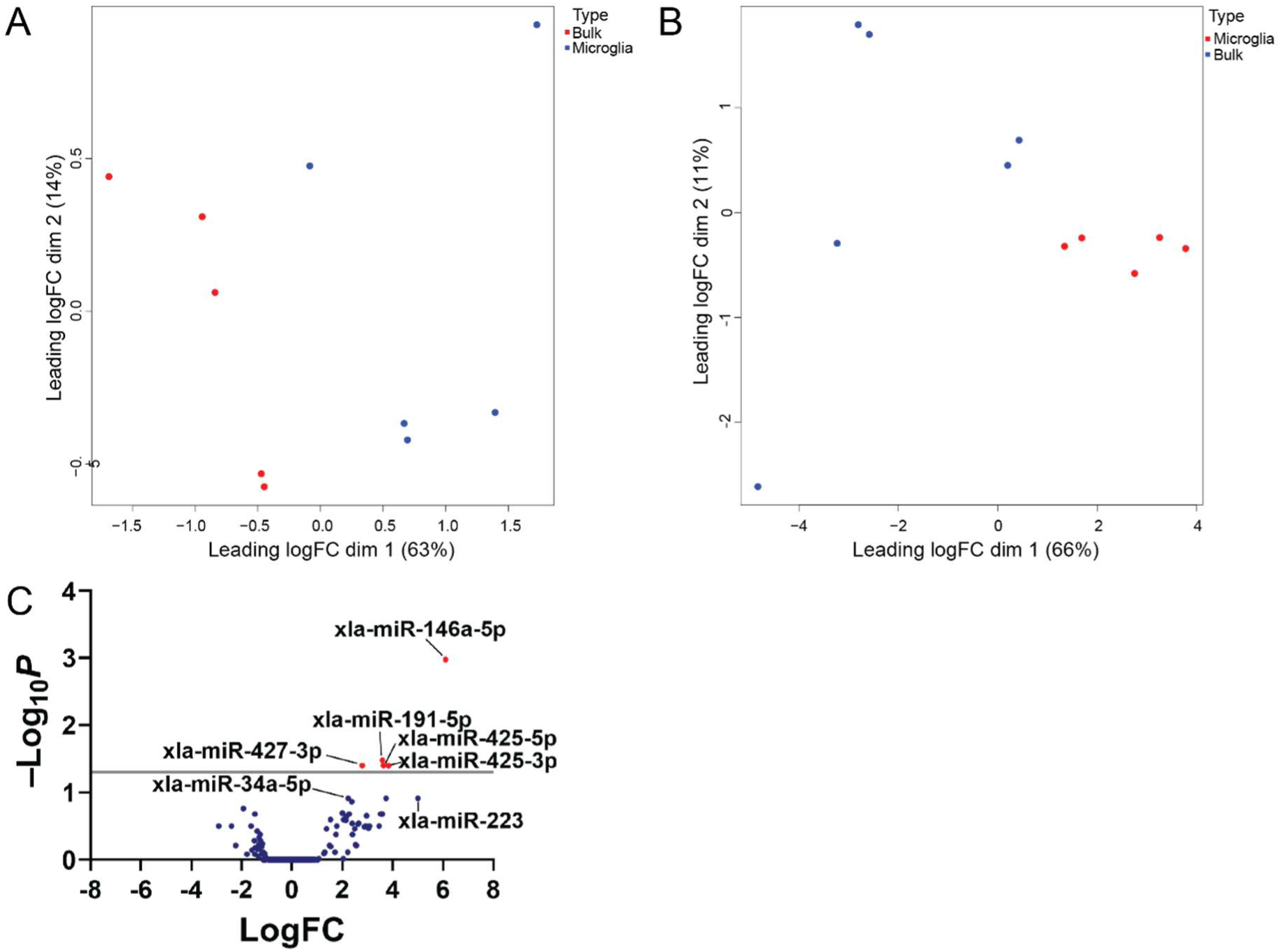
The miRNA transcriptional profile of Xenopus microglial reveals few enriched miRNA compared to non-microglial brain cells. **A.** MDA analysis of miRNA sequencing data from mCherry^+ve^ and mCherry^-ve^ cell types reveals strong clustering by cell type. **B.** MDA analysis of mRNA sequencing data from mCherry^+ve^ and mCherry^-ve^ cell types reveals strong clustering by cell type. **C.** Volcano plot of miRNA that were differentially expressed between pooled mCherry^+ve^ microglia and mCherry^-ve^ bulk brain cells. Six of 213 miRNA were enriched in mCherry^+ve^ microglia compared to mCherry^-ve^ bulk brain cells (log2FC > 1, adj.P.Val <0.05). N= 5 mCherry^+ve^, 5 mCherry^-ve^.

A total of 213 miRNA transcripts, and 25,534 mRNA transcripts were identified (Fig.3C). Amongst the 886 mRNA transcripts enriched in mCherry^+ve^ microglia were known microglial transcripts including *p2ry12*, *cd86*, and *trem2* (Supplementary Table 4). We identified six miRNA that were enriched in mCherry^+ve^ microglia compared to mCherry^-ve^ bulk brain cells (log2FC > 1, adj.P.Val <0.05). This subset included xla-miR-146a-5p, xla-miR-191-5p, and xla-miR-425-5p, which have previously been reported to be enriched in microglia in other species (Fig.3.C, Supplementary Table 5)^21^. An additional 26 miRNA were enriched but did not reach the threshold for significance after adjusting for covarities the data (log2FC > 1, P.Val <0.05). This group included known microglial miRNA such as xla-miR-223-3p and xla-miR-34a-5p (Fig.3.C)^21^. The lack of statistical significance is likely due to the limited sample size available for this study. The sequencing results derived from this study were taken forward into a cross-species analysis to identify a conserved microglial miRNA signature as well as the identification of miRNAs involved in demyelination and remyelination.

### Cross species comparison reveals a conserved microglial miRNA signature

Identifying miRNA orthologs with conserved enrichment can reveal key regulators of the microglial lineage and provide a framework for using preclinical models to study human-relevant miRNAs. To identify conserved microglial miRNAs, we compared our human and *Xenopus* miRNA datasets with a mouse microglial miRNA dataset we previously generated^21^. We identified 90 miRNA that were expressed by microglia in all three species (Fig. 4A), and four of these showed conserved enrichment compared to their respective non-microglial bulk samples (adj.P <0.05) (Fig.4B, Supplementary Table 6). These miRNA (miR-146a-5p, miR-191-5p, miR-34a-5p, miR-425-5p) have previously been identified as regulators of microglial functions or as miRNA with a key role in the pathogenesis of neurodegenerative diseases, including MS^21,28,37,38^. The conservation of these miRNAs across species separated by >350 million years of evolutions highlights their integral role in establishing and maintaining microglial identity and function^15^.

**Figure 4.**
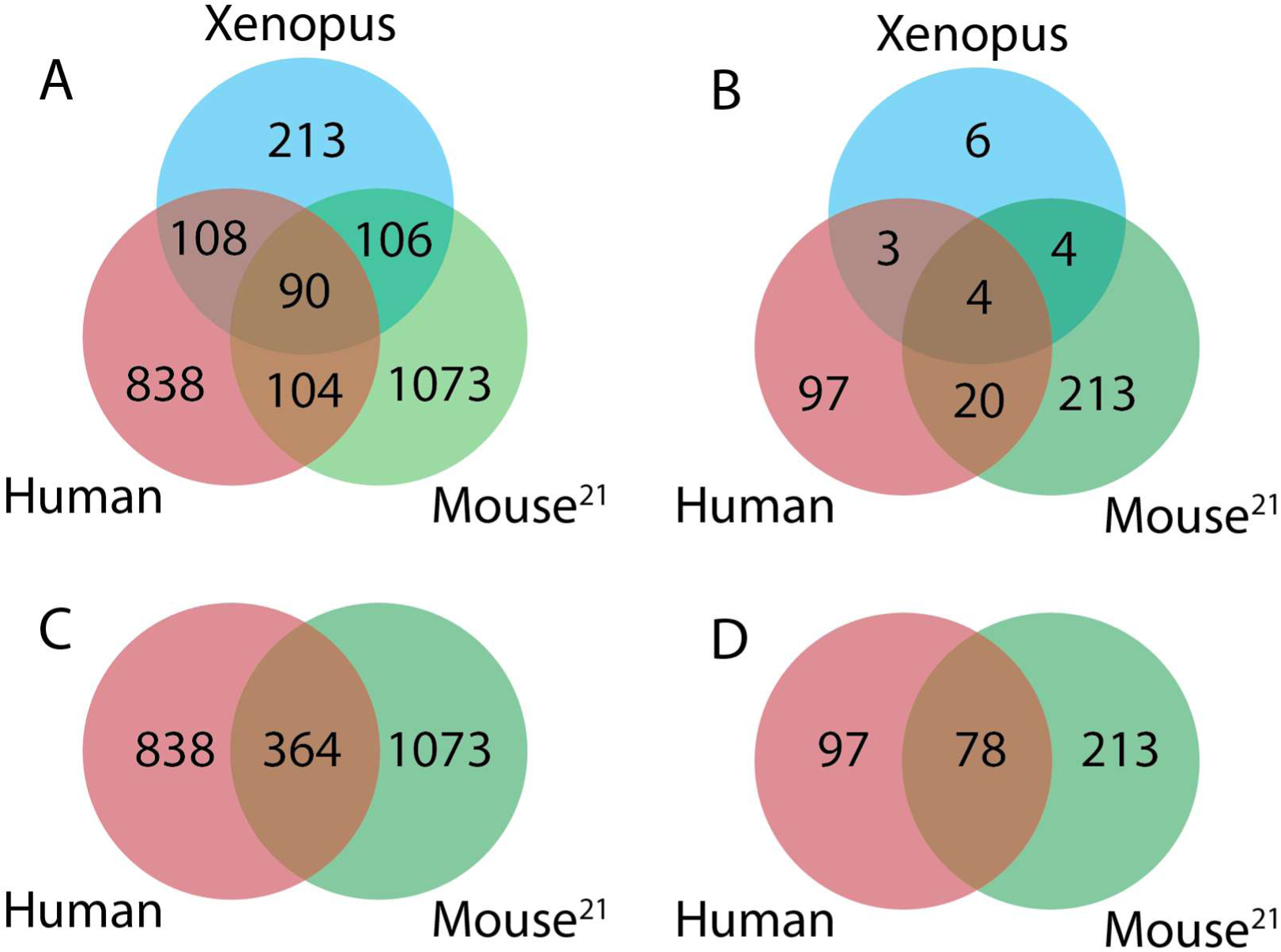
Cross species analysis of microglial miRNA expression and enrichment reveals conservation of a small set of miRNA across three species. **A.** Venn diagram comparing miRNA expression between human, mouse and xenopus microglia. **B.** Venn diagram comparing miRNA enrichment between human, mouse and xenopus microglia. **C.** Venn diagram comparing miRNA expression between human, mouse and xenopus microglia. **D.** Venn diagram comparing miRNA enrichment between human, mouse and xenopus microglia.

To probe more proximal evolutionary conservation, we ran a secondary analysis restricted to human and mouse data. This analysis identified 364 miRNA orthologs with conserved expression in both human and mouse microglia (Fig.4C). Of these, 78 miRNA orthologs were significantly enriched in both species (adj.P <0.05) (Fig.4D, Supplemental Table 7). This included all four miRNAs identified in the human-mouse-xenopus analysis and many miRNAs previously linked to microglial function^21,39^. Notable examples include established regulators of microglial function, such as miR-223-3p and miR-342-3p, and miR-511-3p which our previous mouse study identified as a likely key regulator of microglial function^21^. Additionally, we identified many miRNAs specifically upregulated in human microglia, including miR-629 which has been implicated in MS pathogenesis^40^.

### Microglial miRNAs are differentially expressed during demyelination and remyelination

Microglia contribute to the pathogenesis of neurodegenerative diseases, including key roles in demyelinating diseases such as MS^9,10^. Previous studies have identified altered brain miRNA expression in people with MS and in animal models of myelin damage, such as the cuprizone model^28–30,37,41,42^. Therefore, we investigated the dynamic expression of microglial miRNA in the cuprizone model. We purified CD45^+ve^ microglia from the brains of mice at 5-weeks of cuprizone induced demyelination and 1-week of remyelination using immunopanning and sequenced the miRNAome (Fig.5A).

**Figure 5.**
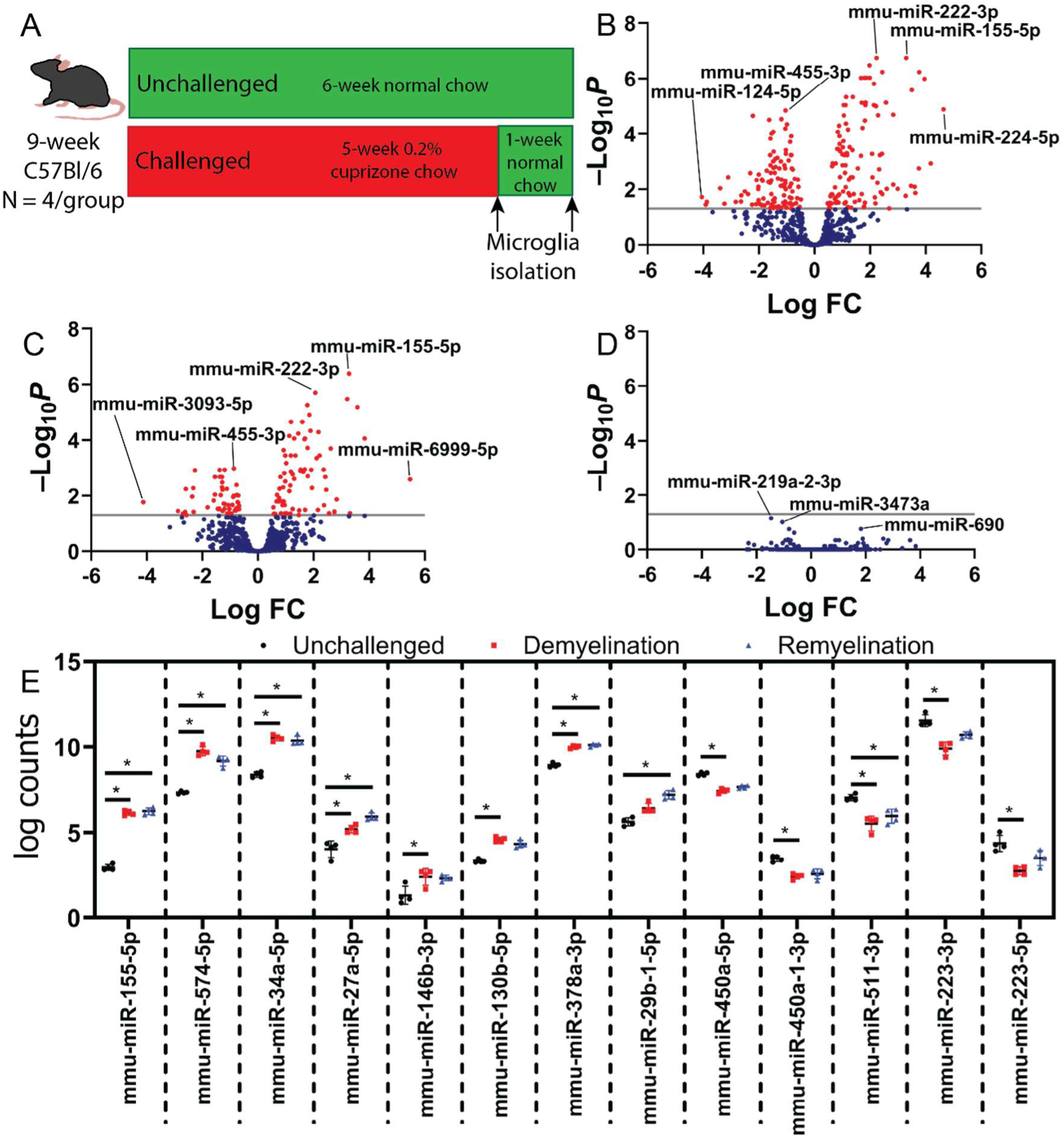
Mouse microglia differentially express miRNA during demyelination and remyelination. **A.** Schematic showing the experimental design of the cuprizone experiment. **B.** Volcano plot of miRNA that were differentially expressed between pooled CD45^+ve^ microglia from unchallenged and demyelinated mouse brains. Out of 700 miRNA 86 were downregulated and 83 were upregulated in CD45^+ve^ microglia from demyelinated (5 weeks) mice compared to CD45^+ve^ microglia from age matched unchallenged mice (log2FC > 1, adj.P.Val <0.05). **C.** Volcano plot of miRNA that were differentially expressed between pooled CD45^+ve^ microglia from unchallenged and remyelinated mouse brains. Out of 700 miRNA 37 were downregulated and 52 were upregulated in CD45^+ve^ microglia from remyelinated (1 week) mice compared to CD45^+ve^ microglia from age matched unchallenged mice (log2FC > 1, adj.P.Val <0.05). **D.** Volcano plot of miRNA shows that no miRNAs were differentially expressed between pooled CD45^+ve^ microglia from demyelinated and remyelinated mouse brains (log2FC > 1, adj.P.Val <0.05). **E.** Expression data of 13 miRNAs that showed conserved enrichment in human and mouse microglia and were differentially expressed during demyelination and/or remyelination. Green background indicates miRNA that also showed conserved enrichment in xenopus. N= 4 mice/group.

We identified 700 miRNA, including 641 miRNA we had previously identified in developmental mouse microglia dataset^21^. The 48 miRNAs that were unique to this study had relatively low expression in microglia from unchallenged mice (log CPM -1.097±1.265, mean±SD), likely explaining why they did not reach the threshold for detection previously. Of these 48 miRNAs, 10 were upregulated, in microglia during demyelination and/or remyelination compared to age matched controls, (log2FC >±1, adj.P.Val <0.05); however, only two of these 10 have identified human orthologs (miR-155-3p and miR-224-3p). The remaining 38 miRNAs were not significantly changed. During demyelination, 86 miRNAs were downregulated and 83 were upregulated, while during remyelination 37 miRNAs were downregulated and 52 were upregulated relative to age matched controls (log2FC >±1, adj.P.Val <0.05) (Fig.5B, C; Supplementary Tables 8 and 9). No miRNAs were differentially expressed when comparing microglia from remyelination and demyelination samples (Fig.5D).

We next analysed our datasets to identify miRNAs that are both enriched in microglia across multiple species and are also dysregulated in the cuprizone model, as these miRNAs are likely to have key roles in regulating microglial function and potentially in myelin repair. All four miRNAs orthologs that were identified in the human-mouse-xenopus dataset were expressed in the cuprizone dataset, but only miR-34a-5p showed differential expression during demyelination or remyelination. In comparison, all 78 miRNAs orthologs that were identified in the human-mouse dataset were expressed in the cuprizone dataset. Among the differentially expressed miRNAs, 13 showed conserved enrichment in microglia in both humans and mice (Fig.5E), 44 miRNAs were enriched in mouse but not human microglia, and 153 miRNAs were not enriched in human microglia or in microglia from unchallenged mice^21^. Of the 13 conserved differentially expressed miRNAs, eight have orthologs previously identified as differentially expressed in the brains of people with MS (miR-155-5p, miR-574-5p, miR-34a-5p, miR-27a-5p, miR-146b-3p, miR-130b-5p, miR-223-3p, miR-223-5p), while five have not previously been reported in MS (miR-378a-3p, miR-29b-1-5p, miR-450a-5p, miR-450a-1-3p, miR-511-3p; Fig.5E)^28–30,37,41,43,44^. While many of these miRNAs have been previously implicated in MS, studies demonstrating cell specific expression in the brains of people with MS have been lacking, with the notable exception of miR-155-5p^44^. Our findings highlight human-relevant miRNAs with potential roles in demyelination and remyelination.

Many of the miRNAs differentially expressed in microglia during the cuprizone model were not enriched in human microglia. These included miRNAs both with and without human orthologs and may therefore also contribute to microglial responses in neurological disease. miRNAs that were not enriched in human or mouse microglia accounted for many of the most strongly upregulated transcripts, representing the 10 and nine most up regulated miRNA during demyelination and remyelination respectively, compared to microglia from unchallenged mice. The most strongly upregulated miRNA with a human ortholog in both demyelination and remyelination was mmu-miR-224-5p (4.8 and 3.8log fold increases respectively). This miRNA has been reported to have roles in macrophage function in osteoarthritis and is elevated in the plasma of people with amyotrophic lateral sclerosis^45,46^. Among miRNAs with a human ortholog that were downregulated compared to control, mmu-miR124-5p was the most strongly downregulated during demyelination (-4.0 log fold) and the second most strongly downregulated during remyelination (-2.7 log fold). miR-199a-5p was the most strongly downregulated during remyelination (-2.7 log fold). In addition, miR-124-3p was also downregulated compared to unchallenged microglia at both timepoints (-1.5 and -1.3 log fold change respectively). miR-124 is among the most studied miRNA in microglial biology, however, its key role in neuronal function raises concerns about its clinical utility for modulating microglial activity^47^. In contrast, miR-199a-5p has been identified as a potential serum biomarker in people with MS, its downregulation in this study suggests potential divergent function in the CNS vs peripheral cells^48^. These data demonstrate the importance of these miRNAs in microglia during demyelination and remyelination and identify additional miRNA with potential clinical relevance.

### miR-511-3p regulates efferocytosis of myelin debris

According to miRbase, miR-511-3p orthologues exist in several mammalian species including mice, humans, rats, and rhesus monkey but are absent in non-mammalian species including *X. laevis* (miRbase access date 29/10/2025). Across our datasets miR-511-3p is consistently enriched in microglia in species where it is expressed (Fig.6A,B). Additionally, microglial mmu-miR-511-3p expression is significantly reduced at 5-weeks demyelination and 1-week remyelination in the cuprizone model (Fig.5E). Together, these finding suggest that miR-511-3p may have an important role in microglial identity and function. miR-511-3p has previously been implicated in inflammation and polarisation of peripheral macrophage populations^49–51^. In both mice and humans, miR-511-3p is an intronic miRNA located within the mannose receptor gene (*MRC1*), with which it is transcriptionally co-regulated^51^. *MCR1* encodes a C-type lectin involved in the cellular uptake of endogenous and exogenous ligands^52^. Based on these observations, we investigated if miR-511-3p regulates efferocytosis of myelin debris in human and mouse microglia.

**Figure 6.**
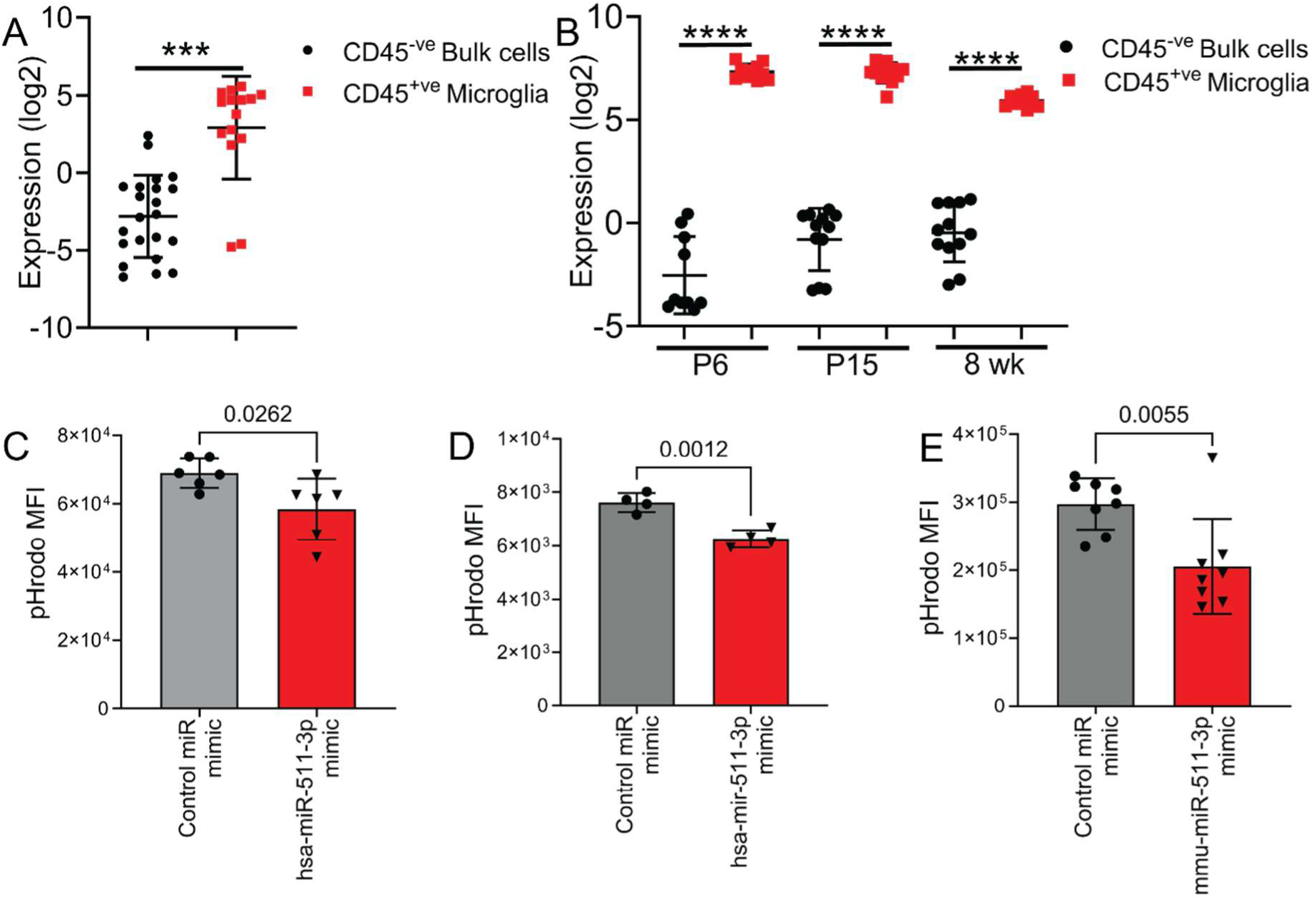
miR-511-3p shows conserved enrichment in human and mouse microglia and preservation of function. **(A)** hsa-miR-511-3p is highly enriched in CD45^+ve^ microglia compared to CD45^-ve^ bulk brain cells in human, N= 16 CD45^+ve^, 22 CD45^-ve^. **(B)**. mmu-miR-511-3p is highly enriched in CD45^+ve^ microglia compared to CD45^-ve^ bulk brain cells in human and mouse^21^ N= 10 P6, 12 P15 and 8 week. **(C, D, E)** miR-511-3p regulates efferocytosis of myelin debris. Human iMGLs (**C.** C11, **D.** C107) and mouse primary microglia (**E**) and were transfected with species appropriate miR-511-3p mimics or control mimics. After 24 hours mouse or human GM-CSF (50 ng/ml) was added to cultures for 24 hours to mildly inhibit phagocytosis. Mouse primary microglia and human iMGLs were exposed to pHRodo labelled human myelin debris for 45 minutes and efferocytosis was assessed by flow cytometry. Representative results from three independent experiments. \*\*\**P*<0.001, \*\*\*\**P*<0.0001.

Primary mouse microglia were isolated from mixed glial cultures and human stem cell induced microglial like cells (iMGLs) were generated by differentiation of two human iPSCs cell lines (C11 and C107). Primary mouse microglia and human iMGLs were transfected with species specific miR-511-3p mimics, or control mimics and were exposed to pHRodo labelled myelin debris. Myelin uptake was quantitated by flow cytometry. In both mouse microglia and human iMGLs, cells transfected with miR-511-3p mimics internalised significantly less myelin debris than cells transfected with control mimics, demonstrating a role for miR-511-3p in the regulation of efferocytosis in microglia (Fig 6c, d).

## Discussion

Microglia, the brains resident immune cells, are intrinsic to the development and homeostasis of the CNS and are also central to CNS inflammatory responses and the pathogenesis of neurological diseases^4–10^. There is much interest in the regulation of microglial function as a treatment for neurodegenerative diseases such as MS. An understanding of the factors that regulate microglial gene expression will likely be crucial to achieving this goal. miRNAs are one of the most abundant post-transcriptional regulators of gene expression with more than 60% of human protein coding genes being regulated by one or more miRNA^16^. Their critical role in regulating cellular processes indicates the involvement of miRNA in most, and potentially all, developmental, physiological, and pathological processes^17,18^. miRNA loci are found in both intragenic and intergenic regions; ∼50% of vertebrate miRNAs are located in intragenic regions and many of these miRNAs regulate their host genes^15,53^. miRNAs have a deep evolutionary history and expansions in the miRNA repertoire coincides with major evolutionary events including development of complex bilaterians, and the origins of vertebrates, placental mammals, and primates suggesting miRNA may have significantly contributed to these evolutionary events^15^. The largest expansion of intragenic miRNA coincides with the advent of neural crest development with 21 new miRNA families emerging within the introns of genes that regulate neuronal development^15^. This shared evolutionary history further highlights the critical role of miRNA in the regulation of gene expression and suggests gene expression cannot be fully understood independent of miRNA expression. Insights into microglial miRNA and mRNA have been found in mouse studies^20,21,24^. However, while the mRNA transcriptome of human microglia has recently been investigated^22,54,55^ studies of miRNA in human microglia have been limited.

In this study we undertook a detailed investigation of the miRNAome of human microglia and identified a core miRNA signature in microglia. We further identified the expression of miRNA that are conserved between human, mouse, and *X. laevis* microglia, and a set conserved between only human and mouse microglia, which likely represent a subset of miRNA critical to the function and identity of microglia. We furthermore characterised mouse microglial miRNA in the cuprizone model and identified differentially expressed miRNA that may have key roles in the regulation of demyelination are remyelination.

We identified more than 90 miRNA that are enriched in human CD45^+ve^ microglia compared to the CD45^-ve^ bulk brain cells. This finding is consistent with previous studies that have identified unique miRNA profiles in mouse microglia and studies that have consistently identified unique mRNA signatures in human microglia using single cell sequencing^21,24,55–57^. This difference in miRNA signature reflects both the unique function of microglia in the CNS as well as their distinct cellular origin. Microglia are unique amongst the major CNS cell populations, arising from the embryonic yolk sac and entering the CNS during development, whilst other CNS populations (e.g. neurons, astrocytes, oligodendrocytes) are derived from the neuroepithelial lineage^2,3^. The miRNAs most highly enriched in microglia included miRNA previously identified as enriched in mouse microglia (e.g. miR-31-3p and miR-511-3p); but also included miRNA not enriched in mouse microglia (but with mouse orthologs, e.g. miR-653-5p), and miRNAs only reported in humans (e.g. miR-4645-3p and miR-5193).

In mice, studies have shown that expression of miRNA and mRNA is influenced by multiple factors including age, sex, and brain region^21,24,58,59^. In this study we isolated microglia from fresh brain tissue from non-lesion areas of surgical resections for intractable epilepsy. These microglia expressed high levels of mRNA associated with a homeostatic phenotype such as *P2RY12* and *CX3CR1*^54^. Consistent with our findings in mice^21^ we did not observe that sex significantly affected human microglial miRNA expression. Additionally, we did not find that brain location or specific form of epilepsy influenced miRNA expression.

As miRNAs are a highly conserved mechanism for post translational regulation of gene expression, we hypothesised that microglia would have cross species overlap in miRNA expression, with miRNA that are upregulated across species representing miRNA with critical roles in microglial function. To assess this, we compared our human microglia miRNA data with microglia expression in mice and *X. laevis*, two common laboratory models used to study the CNS. Unlike the human microglial miRNA dataset and the moue microglial miRNA dataset^21^ relatively few miRNAs (six miRNA) were found to be enriched in mCherry^+ve^ microglia compared to mCherry^-ve^ bulk brain cells, although a further 32 were enriched but did not reach statistical significance after adjusting for covariates.

Comparison of miRNA enriched in microglia in humans, mice, and xenopus identified only four miRNAs with conserved microglial enrichment. Conversely, comparison of enriched microglial miRNA in only humans and mice revealed 78 conserved miRNAs. The four miRNAs with conserved microglial enrichment in all three species (miR-146a-5p, miR-191-5p, miR-34a-5p, miR-425-5p) have previously been identified as highly enriched microglial miRNA^21^. Furthermore, these miRNAs have been implicated in CNS disease including MS. miR-191 was found to be significantly downregulated in normal appearing white matter^37^, while miR-34a and miR-146a are significantly upregulated in CNS lesions in people with MS^28^. miR-425 has been implicated in Parkinson’s disease^60^ and high levels of serum miR-425 was associated with increased white matter lesions in people with MS^61^.

Large expansions in miRNA number occurred during important evolutionary events including at the advent of vertebrates (human, mouse, *X. laevis*), placental mammals (human, mouse), and primates (human)^15^. Further diversification of miRNAs was significantly influenced by whole genome duplication and the subsequent deletion of redundant genes. The conservation of microglial enrichment of four miRNAs over >350 million years suggested the key role of these miRNAs in microglial identity and function^15^. However, there has also been clear divergence of microglial miRNA expression between species. While this has not been previously investigated it has been reported that the distinct morphology of ramified homeostatic microglia is conserved over more than 400 million years of evolution as is a core microglial transcriptional signature^1^. It is interesting that in this study the transcriptional clustering of microglia aligns with expansions of miRNA; non-mammalian microglia (zebrafish and chicken) were transcriptionally the most distinct from humans which clustered most strongly with other mammals, particularly other primates^1^. As intronic miRNA share an evolutionary history with their host gene^15^, understanding miRNA profiles may be critical to fully understanding transcriptional states in many cell types including microglia.

miRNAs have also been identified as dysregulated in CNS disease including MS^28–30^. We investigated microglia miRNA expression in the cuprizone model to understand the role of microglial miRNA in demyelination and remyelination. Knockout of all miRNAs in microglia via microglia specific dicer deletion results in enhanced demyelination are impaired remyelination, however the involvement of specific miRNA in this phenotype has not been determined^25^. We identified dysregulated miRNA during demyelination (86 downregulated and 83 upregulated) and remyelination (37 miRNAs downregulated 52 upregulated) compared with age matched controls. Unexpectedly, we did not identify differences between microglial miRNA expression between demyelination and remyelination timepoints. Microglial play multiple roles in the cuprizone model including the clearance of myelin debris and the promotion of remyelination^32,33,62^. In the cuprizone model oligodendrocyte death is evident from 2-weeks cuprizone administration which coincides with the beginning of microglial accumulation in the corpus callosum^63–65^. Microglial continue to accumulate in the corpus callosum until 5-weeks which corresponds to peak demyelination^63–65^. Oligodendrocyte progenitor cell (OPC) proliferation and remyelination occurs concomitantly with late demyelination and continues following cuprizone removal^63–65^. Thus, it is possible that the selected timepoints (5-weeks demyelination and 1-week remyelination) do not represent fully distinct microglial states but instead microglia transiting between states associated with damage and repair. Assessment of the microglial miRNAome in earlier stages of the cuprizone model may further enhance our understanding of the microglial miRNA that regulate clearance of myelin debris, while this current study focuses on microglial miRNA in late-stage demyelination and remyelination.

Of the miRNAs linked to microglial function miR-155 and miR-124 have been most consistently connected to neuroinflammatory and anti-inflammatory microglia respectively^39^. In this study miR-155-5p and miR-124-5p are among the most strongly differentially expressed microglial miRNAs during both demyelination and remyelination, with miR-155-5p upregulated and miR-124-5p down regulated at both timepoints. miR-155-5p is enriched in both mouse and human microglia and its expression is the cuprizone model is consistent with reports of miR-155-5p as a key promotor of microglial inflammation that is elevated in MS brain tissue and MS animal models^28,39,41,42,44^. The role of miR-124 in microglia MS is more complex. miR-124 is one of the most abundant miRNAs in the CNS with miR-124 transcripts representing ∼25-50% of miRNA in the mouse CNS and is highly enriched in mouse microglia compared to peripheral macrophages^66,67^. However, we did not identify miR-124-5p as an enriched miRNA in either humans or mice compared to bulk brain cells, consistent with its reported high expression in neurons and role in neurogenesis and neuronal function^47^. miR-124 is upregulated in hippocampal neurons in MS brain tissue and mouse cuprizone tissue, and expression is associated with demyelination suggesting a potential pathogenic role for miR-124 in MS^68^. Conversely, we found that mmu-miR-124-5p expression was significantly reduced in microglia during both demyelination and remyelination compared to unchallenged control microglia. Furthermore, miR-124-5p has been reported to inhibit OPC differentiation suggesting that lower expression of miR-124-5p in OPC may be beneficial for myelination^69^. Together these data suggest different roles of miR-124 in different cell types and highlight the importance of considering miRNA expression and regulation on a cellular level when considering the use of miRNA for therapeutic development.

We identified significant conservation of microglial miRNA enrichment between humans and mice, however, each species also expressed distinct miRNAs including both orthologous and non-orthologous miRNAs. We sought to identify the conserved enriched microglial miRNA in these datasets as these likely have key roles in regulating microglial function and will have relevance to humans while possessing the capacity to be studied in preclinical models. Of the 78 miRNAs enrichment conserved in microglia between mice and humans 13 were differentially expressed during demyelination and or remyelination in the cuprizone model (8 increased and 5 decreased in at least one timepoint compared to unchallenged mice). Several studies have investigated miRNA in the brains of people with MS; however, these data come from bulk tissue studies and thus it is unclear the extent to which microglial miRNAs contribute to this signature^28–30,37,41,43,68^. Across these studies no miRNA has been consistently reported as deregulated; for example, miR-155 has been identified as both increased^28,41,43^ and unchanged^29^ in white matter lesions, while other studies did not report on miR-155 expression in grey matter lesions^30^ and normal appearing white matter^37^. Although variation in sample processing and miRNA identification techniques likely contributes to this apparent discrepancy a key difference in these studies is the location and type of brain tissue analysed (active and inactive white matter lesions^28^, white matter lesion^29^, grey matter lesions^30,41^, hippocampal lesions^68^, and normal appearing white matter^37,43^). miRNA has been shown to have distinct tissue specific profiles in both humans and mice^66,70^ and within tissues cell types, including microglia^21^, also have distinct miRNA profiles. As different lesion types have different cellular compositions^71^ the identification of differentially expressed miRNA will likely be significantly influenced by lesion type. Thus, to fully understand the role of dysregulated miRNA in CNS disease it is critical to understand the cell specific expression of miRNA.

Overall, we found significant overlap in the expression of miRNA in the brains of people with MS and in mouse microglia during demyelination and remyelination. The 13 orthologous miRNAs identified in this study may represent miRNA that play key roles in MS pathogenesis. These miRNAs included eight that have orthologous that have previously been identified as differentially expression in brain tissue of people with MS^28,29,37,41,43^. Six mouse miRNAs were increased in microglia during demyelination and/or remyelination in cuprizone have orthologs that have been found to be increased in MS brain samples in at least one study (mmu-miR-155-5p, mmu-miR-574-5p, mmu-miR-34a-5p, mmu-miR-27a-5p, mmu-miR-146b-3p, and mmu-miR-130b-5p)^28,29,41,43^. Two mouse miRNAs were decreased in microglia demyelination in cuprizone but have been found to be increased in MS brain samples (mmu-miR-223-3p and mmu-miR-223-5p)^28,37^.

Our study identified several miRNA that have not previously been identified as microglial regulators in MS or its animal models including five miRNA with orthologues that show conserved microglial enrichment between humans and mice (mmu-miR-378a-3p, mmu-miR-29b-1-5p, mmu-miR-450a-5p, mmu-miR-450a-1-3p, and mmu-miR-511-3p). It is possible that these miRNAs may be differentially expressed specifically in microglia in MS but could not be detected due to dilution of signal due to other cell types. These miRNAs may represent novel miRNA targets to regulate microglial function in neurodegenerative and demyelinating diseases. We further investigated the function of miR-511-3p in microglia due to its role macrophage function and activation^49–51^. miR-511-3p is highly expressed in macrophages activated into anti-inflammatory like states including “M2” macrophages and tumour associated macrophages (TAMs)^49,50^. We found that elevation of intracellular miR-511-3p levels resulted in decreased uptake of myelin debris by both human iMGLs and primary mouse microglia. Uptake of myelin debris by microglia is vital to efficient remyelination^62^. The reduced expression of microglia mmu-miR-511-3p expression in the cuprizone model may be related to the need to upregulate myelin clearance pathways in response to demyelination. miR-511-3p is located within the MRC1 gene in both mice and humans which encodes the mannose receptor (MR), miRNAs frequently regulate their host gene and expression of miR-511-3p is higher in TAMs with elevated MR expression suggesting co-expression^49,51^. The MR is a C-type lectin that functions as an endocytic receptor for endogenous and pathogen associated molecules^52^. Myelin debris can be taken up through multiple pathways including endocytosis^72^, this it is likely that regulation of MR expression is one mechanism through which miR-511-3p regulates myelin debris uptake. These data identify a targetable pathway to promote myelin debris clearance in MS to enhance remyelination.

In summary we investigated the miRNAome of microglia in three species, humans, mouse, and *X. laevis*, to identify the conserved miRNAs that are important to the identity and function of microglia. We identified subsets of miRNAs with conserved microglial enrichment across all three species as well as those with conserved enrichment only between humans and mice, which are likely important to microglial identity. Differential expression of miRNAs plays a major role in neurological diseases; thus, we further investigated microglia miRNAs in the cuprizone model and identified conserved microglial enriched miRNAs that were differentially expressed during demyelination and remyelination. We further investigated the function of one of these miRNAs, miR-511-3p in microglial and identified that elevated miR-511-3p impaired myelin debris uptake in both primary mouse microglia and human iMGLs. The miRNA identified in this study are potential therapeutic targets for the regulation of microglial function in neurological disease.

## Methods

### Animals

C57Bl/6 mice were obtained from Ozgene (Perth, WA, Australia). Mice were maintained in 12-hour light/dark cycle and were provided with food and water ad libitum. Mouse experiments were approved by the Florey Institute of Neuroscience and Mental Health Animal Ethics Committee and were performed according to National Health and Medical Research Council (Australia) guidelines.

*Xenopus laevis* [*Tg*(*MPEG-mcherry)*] were at the Institut de Cerveau (ICM, Paris Brain Institute), Paris, France^35^. All experiments used development stage 46–49 tadpoles according to the Nieuwkoop and Faber (NF) stages of *Xenopus laevis* development^73^ and were performed in accordance with institutional and national guidelines.

### Human Sample demographics and collection

Fresh samples of human brain tissue were collected from 23 individuals who were undergoing surgery to remove lesions associated with intractable focal epilepsy. The tissue samples were from normal appearing tissue removed to allow access to the epileptic lesion. Written informed consent was obtained from all donors over the age of 18, or from their legal guardians for donors under the age of 18. This research was designed in accordance with national standards as outlined in the National Statement on Ethical Conduct in Human Research (Australia) (https://www.nhmrc.gov.au/guidelines-publications/e72) and was approved by the Research Ethics Committee. Details of the human samples can be found in Supplementary Table 1.

### The cuprizone model of demyelination and remyelination

C57Bl/6 mice aged 9 weeks were fed 0.2% cuprizone (w/w) cuprizone (bis-cyclohexanone oxaldihydrazone, Merck, Darmstadt, Germany) in powdered chow (Barastoc, Pakenham, Victoria, Australia) for 5 weeks. Cuprizone chow was changed daily, and mice were housed in pairs to ensure adequate consumption. To induce remyelination cuprizone treated mice were fed standard chow for one week. Age-matched control mice were fed standard chow throughout. Brains were collected at 5-weeks demyelination (14 weeks old) or one-week remyelination (15 weeks old). Brains from control mice were collected at 14-weeks old. The meninges, olfactory bulb, and cerebellum were removed, and microglia were purified by immunopanning.

### Microglial purification by immunopanning and RNA extraction from mouse and human brain tissue

CD45^+ve^ microglia and CD45^-ve^ bulk brain cells were purified from human and mouse brain tissue by immunopanning as previously described^21^. Briefly, immunopanning plates (one per sample) were prepared by coating a 10 cm petri dish with goat anti-rat IgG (3 µl/ml in 50 mM Tris-HCL pH 9, 112-005-167 Jackson Immunoresearch, PA, USA) and incubating overnight at 4°C. Plates were washed 3 times with DPBS (Ca+/Mg+, Merck) and incubated with anti-CD45 antibody (1.67 µl/ml 12 ml 0.2% BSA in DPBS (Ca+/Mg+), 550539, BD Biosciences, NY, USA) for at least 2 hours prior to use isolating microglia.

Brain tissue was minced into ∼1 mm^3^ pieces and incubated in papain buffer (20 U/ml papain and 12,500 U/ml DNase I (Worthington, OH, USA), 0.46% glucose, 26 mM NaHCO3 0.5 mM EDTA, 1x EBSS) at 34°C for 100 minutes with a constant carbogen flow and periodic shaking. Tissue was then washed in low ovomucoid solution to inhibit the remaining papain (0.015 g/ml trypsin inhibitor (Worthington) in DPBS (Ca+/Mg+) at pH of 7.4), followed by gentle trituration in low ovomucoid solution to generate a single cell suspension. High ovomucoid solution (0.030 g/ml BSA (Merck), 0.030 g/ml trypsin inhibitor (Worthington) in DPBS at pH of 7.4) was layered underneath the cell suspension and cells were centrifuged at 250 g for 15 minutes. Cells were resuspended in 0.02% BSA in DPBS (Ca+/Mg+), transferred to anti-CD45 coated immunopanning plates, and incubated at room temperature for 30 minutes with periodic agitation. Unbound cells CD45^-ve^ bulk brain cells were collected, centrifuged at 300 g for 5 minutes, and resuspended in 700 µl Qiazol (Qiagen, Venlo, The Netherlands). The panning plates were washed 3x with DPBS (Ca+/Mg+) following which 700 µl of Qiazol (Qiagen) was added and the plates were scraped to collect the bound cells C45^+ve^ microglia. Total RNA was extracted using an miRNeasy mini kit (Qiagen) according to manufacturer’s instructions.

### Microglial purification by FACS and RNA extraction from X. laevis brains

Stage 46–49 *Tg*(*MPEG-mcherry) X. laevis* tadpoles were euthanized in 0.5% MS-222 (ethyl-3-aminobenzoate methanesulphonate; Cayman Chemical) and brains were collected into Hank’s balanced salt solution (HBSS) without calcium and magnesium, (ThermoFisher MA, USA, 141175-095). Brains from 30–40 tadpoles of either sex were pooled per sample and were digested in dissociation media containing collagenase D (0.05%, Roche, Basel, Switzerland, 1-088-874), dispase (0.5%, Roche, 165859), DNase I (0.025 U/ml, Sigma-Aldrich, MA, United States, D4527) and N-alpha-tosyl-L-lysine chloromethyl ketone hydrochloride (0.1 µg/ml, Sigma-Aldrich, cat. no. T7254) for 15 minutes at room temperature prior to mechanical dissociation. Cells were washed in HBSS and FACS was used to separate *MPEG-mcherry^+^* microglia from non-microglial brain cells. Microglia and non-microglia brain cells were resuspended in 700 μl of Qiazol (Qiagen) and total RNA was extracted using an miRNeasy mini kit (Qiagen) according to manufacturer’s instructions.

### miRNA and mRNA library construction and sequencing

RNA quality was evaluated using a bioanalyser or Qubit and TapeStation, all samples were found to have an RNA integrity score (RIN) greater than 7 and were used for library construction. Human small RNA library construction and sequencing were performed at Micromon Genomics (Monash University, Melbourne, Australia). Briefly, small RNA libraries were prepared using the NEXTFlex Small RNA chemistry V3 (Perkin- Elmer, CT, United States). Libraries were pooled and sequenced with MGISEQ2000-RS (BGI, Shenzhen, Guangdong, China) sequencing technology (single-end 50bp). Mouse and *X. laevis* small RNA library construction and sequencing were performed at Victorian Clinical Genetics Services (Murdoch Children’s Research Institute, Parkville, Australia). Briefly, small RNA libraries were prepared using NEXTFLEX small RNA v4 gel-free kit (Perkin- Elmer). Libraries were pooled and sequenced using NovaSeq 6000 (Perkin- Elmer) sequencing technology (paired-end 2x50 bp).

mRNA library construction and sequencing were performed at Victorian Clinical Genetics Services for all species (Murdoch Children’s Research Institute). mRNA libraries were prepared using Illumina stranded mRNA kit (CA, United States) and sequenced with NovaSeq 6000 (Perkin-Elmer) sequencing technology (paired-end 2x150 bp).

### Processing and analysis of sequencing data

We removed the small RNA adapter sequences and filtered the remaining reads by size (>18 bp) using fastp (ver 0.20.0). miRDeep2 (default settings, ver 0.1.3) was used to map the trimmed reads to species specific reference genomes (*H. sapiens* GrCh38/hg38, *M. musculus* mm10/GRCm38, *X. laevis* Xenopus_laevis_v10.1). To generate gene read counts we utilised the featureCounts pipeline from Subread (ver 2.0.0) with the following flags (-T 8 -s 2 -p -a).

We utilised limma (ver 3.58.1, 10.18129/B9.bioc.limma) and edgeR (ver 4.0.16, 10.18129/B9.bioc.edgeR) for normalisation and differential expression analysis of miRNA^74^. Filtering of genes was performed using the filterByExpr function from the edgeR package and normalisation of gene counts was performed using the Trimmed mean of M values (TMM, edgeR) normalisation method in conjunction with voomWithQualityWeights (limma). Differential expression analyses were used to assess miRNA and mRNA that are enriched in microglia by direct comparison of miRNA or mRNA expression between all CD45^+ve^ microglia and all CD45^-ve^ bulk samples using the TREAT method with a minimum threshold of a ± 2-fold change (Log2FC ± 1) in expression with an FDR < 0.05^75^. The effect of cell type, age, brain region, sex, and disease diagnosis on gene expression variance was assessed by Multi-Dimensional Scaling (MDS) analysis and linear regression modelling. Square root normalisation was used for age analysis as younger age was more frequent in the donors and the miRNA expression data was fitted with a linear mixed model to measure the change in expression over time (years). MDS plots were generated with ggplot2 (ver 3.5.2 (https://cran.r-project.org/web/packages/ggplot2/index.html).

### Determination of cellular proportions

The cellular proportions in human CD45^+^ and CD45^-^populations were determined using the CIBERSORTx High-Resolution Docker container^76^. Reference gene expression matrices for microglia, oligodendrocytes, OPCs, astrocytes, excitatory neurons, and inhibitory neurons were generated from published single-nucleus RNA sequencing^31^. Gene expression matrices were generated for human CD45^+^ and CD45^-^populations and the reference were used to identify cellular proportions using CIBERSORTx with ‘Smode’ batch correction, faction set to 0.25, and sampling set to 1.

### Cross species analysis of miRNA expression and enrichment

Mature miRNA sequences for mmu, hsa and xla were downloaded from miRbase. Blastn was run on all pairwise combinations of miRNA sequences. To identify conserved orthologs between species, blast results were filtered for an e-value < 0.01. Sequences were then filtered for matching miRNA family names (eg let-7, miR-124). To identify conserved enrichment of orthologous miRNAs, independent limma-voom differential expression analyses for CD45^+ve^ microglia vs CD45^-ve^ Bulk in each species (logFC > 0, adj.P < 0.05) for miRNA orthologs were directly compared. Moderated t-statistics are reported for comparison.

### Primary mouse microglial culture

Microglia were purified from mixed glial cultures as previously described^5^. Briefly, brains were removed from postnatal day (P) 0–3 C57Bl/6 mice and the cortical sheets were isolated. Cortices were minced to ∼1mm^3^ and incubated in 0.15 mg/mL trypsin (Merck) in digestion buffer (8 U/ml DNase I (Merck), 1 mM sodium pyruvate, 10 mM HEPES, 1.375 mg/ml glucose, 1.164 mM MgSO4, 3 mg/ml BSA in HBSS) for 15 minutes at room temp and an equal volume of wash solution was added (1.36 U/ml DNase I, 90 µg/ml trypsin inhibitor (Merck), 1 mM sodium pyruvate, 10 mM HEPES, 1.375 mg/ml glucose, 1.419 mM MgSO4, 3 mg/ml BSA in HBSS). The tissue was centrifuged at 300 g for 5 minutes and resuspended in trituration buffer (8 U/ml DNase I, 530 µg/ml trypsin inhibitor, 1 mM sodium pyruvate, 10 mM HEPES, 1.375 mg/ml glucose, 2.664 mM MgSO4, 3 mg/ml BSA in HBSS). Tissue was gently triturated through a 5 ml serological pipette and then a 22g needle to generate a single cell suspension and centrifuged at 300 g for 5 minutes. Cells were resuspended in full DMEM (5% (v/v) heat inactivated foetal bovine serum (Bovogen Biologicals, Melbourne, VIC, Australia), 0.4 mM GlutaMax, 5 U/ml penicillin/streptomycin, 110 mg/ml sodium pyruvate, 5 μg/ml insulin, and 25 mM HEPES in DMEM, ThermoFisher) and seeded in poly-D-lysine (PDL, Sigma-Aldrich) coated flasks at 1-1.4 brains per flask and cells were cultured at 37C/5%CO2. Media was changed the next day and every 3 days after. Cells were collected at 10–14 days in culture depending on microglial density through differential adherence. Purified microglia were centrifuged at 300 g for 5 minutes and resuspended in microglial growth media (MGM; 1x GlutaMax, 1x penicillin/streptomycin, 100 ng/ml oleic acid (Sapphire Bioscience, Redfern NSW, Australia), 10 ng/ml gondoic acid (Sapphire Bioscience), 1.5 μg/ml cholesterol (Avanti Polar Lipids, Alabaster, AL, USA), 10 ng/ml murine M-CSF (PeproTech, Cranbury, NJ, USA), 2 ng/ml human TGF-β2 (PeproTech), 1 μg/ml heparan sulphate (Sapphire Bioscience), 5 μg/ml N-acetylcysteine (Merck), 100 ng/ml sodium selenite (Merck), and 1 mg/ml apo-transferrin (Merck))^77^. Microglia were seeded at 100,000 cells/well in PDL coated 24 well plates and used for downstream assays. Microglia purity was routinely assessed by flow cytometry by labelling with CD11b (BD Biosciences catalogue 564454, 1:100) and consistently found to be >90% pure.

### Human induced microglia-like cell culture

iMGLs were generated from the human induced pluripotent stem cell (iPSC) lines C11 and C107. Prior to induction, iPSCs were routinely maintained in mTESR Plus media (Stemcell Technologies, BC, Canada) in 6 well TC plates (Nunc, NY, USA) coated with Matrigel (Corning, NY, USA). Hematopoietic progenitor cells were first generated from iPSCs using the STEMdiff Hematopoietic Kit (Stemcell Technologies, catalogue 05310) according to manufacturer’s instructions. At the conclusion of the differentiation protocol, hematopoietic progenitor cells were assessed for the surface markers CD34 (Biolegend, CA, USA, catalogue 343507, 1:50), CD45 (BD Biosciences catalogue 563880, 1:50), and CD43 (BD Biosciences catalogue 564543, 1:100), with Zombie NIR™ to assess viability (BioLegend, catalogue 423105, 1:1000) using flow cytometry (CytoFLEX LX, Beckman Coulter, CA, USA). Hematopoietic progenitor cells were only used for differentiation of iMGLs if >90% of cells were CD43+.

iMGLs were generated from hematopoietic progenitor cells using the STEMdiff microglia differentiation kit (Stemcell Technologies, catalogue 100-0019) according to manufacturer’s instructions commencing with 2 x 10^5^ hematopoietic progenitor cells per well of a Matrigel-coated 6 well plate. At the conclusion of the differentiation process, cells were harvested and counted using a haemocytometer. A subset of cells were plated into 24 well plates (Nunc) coated with PDL and collagen for assessment of phagocytosis (∼8 x 10^4^ cells per well) and the remainder replated into Matrigel-coated 6 well plates (∼2 x 10^5^ cells/well) in STEMdiff maturation media (Stemcell Technologies, catalogue 100-0021). At the end of 4 days, cells on Matrigel were harvested and assessed for surface markers CD45 (BD Biosciences catalogue 563880, 1:50), CD11b (BD Biosciences catalogue 564454, 1:20), Trem2 (R&D Systems catalogue FAB17291N, 1:20) and MERTK (R&D Systems catalogue FAB8912A, 1:10), with Zombie NIR™ to assess viability (BioLegend, catalogue 423105,1:1000) using flow cytometry (CytoFLEX LX, Beckman Coulter).

### Transfection of primary mouse microglia and iMGLs

Primary mouse microglia were transfected after 24 hours in monoculture microglia with 100 mM FAM labelled miRNA mimics for mmu-miR-511-3p (Qiagen catalogue 339173, GeneGlobe ID: YM00471881) or a control mimic (Qiagen catalogue 339173, GeneGlobe ID: YM00479902) using TurboFect (ThermoFisher) according to manufacturer’s instructions. iMGLs were transfected after 7 days in STEMdiff maturation media with 100 mM FAM labelled miRNA mimics for hsa-miR-511-3p (Qiagen catalogue 339173, GeneGlobe ID: YM00471883) or a control mimic (Qiagen catalogue 339173, GeneGlobe ID: YM00479902) using TurboFect (ThermoFisher) according to manufacturer’s instructions. After 24 hours media was changed, and cells were used for downstream assays.

### Efferocytosis assay

The efferocytosis activity of mouse microglial and iMGLs was measured by assessing uptake of myelin debris labelled with pHrodo. Human myelin debris were isolated on a Percoll (Merck) gradient and purified as described previously^5^. Myelin debris (100 mg) was incubated with 2 mM pHrodo™ Red (ThermoFisher), succinimidyl ester for 1 hour at room temperature protected from light. The labelled myelin was then washed with sterile PBS and resuspended at 100 mg/ml in sterile PBS. Transfected mouse microglia or iMGLs were incubated with 5 ng/ml GM-CSF (Peprotec) for 24 hours, following which the mouse microglia or iMGLs were cultured with pHRodo labelled myelin for 45 minutes at 37°C/5%CO2. The excess pHrodo labelled myelin was removed and mouse microglia or iMGLs were lifted with trypsin, washed once with PBS and resuspended with PBS with DAPI (1:100). Efferocytosis was assessed using a CytoFLEX LX (iMGLs) or CytoFLEX S (primary mouse microglia) flow cytometer (Beckman Coulter).

### Statistics and software

Analysis of RNA-Seq data was performed as described above. Flow cytometry data was analysed using FlowJo version 10. Efferocytosis data was analysed using GraphPad Prism version 10. Comparisons between two groups were evaluated using a students t-test, P < 0.05 was considered significant.

## Acknowledgments

The authors would like to acknowledge the support of the Australian Government Research Training Program Scholarship (A.D.W.), National Health and Medical Research Council (APP1175775, T.J.K.), Multiple Sclerosis Australia (21-3-038 S.S.; 23-PG-0146, S.S and M.D.B), The Bethlehem Griffis Research Foundation (1906, M.D.B.; 2307, S.S and M.D.B.), and Perpetual Trustees Impact Philanthropy Funding (S.S and M.D.B). This work was partially supported by Inserm, CNRS, Sorbonne-University, the program “Investissements d’Avenir” ANR-10-IAIHU-06 and NeurATRIS, and ANR grant BRECOMY to BZ. The Florey Institute of Neuroscience and Mental Health and the Walter and Eliza Hall Institute of Medical Research acknowledge the strong support from the Victorian Government, particularly the funding from the Operational Infrastructure Support Grant. We thank Dr Jacques Robert (University of Rochester Medical Center, USA) for the generous gift of Tg(mpeg1:mCherry) *X. laevis* transgenic line.

## Author contributions

Conceived and designed the experiments: SS MDB TJK AW BREA BZ. Performed the experiments: ADW SS LBA NSF EM LP MDB. Provision of study materials: PJL, RJL. Analysed and interpreted the data: ADW SS BREA MDB. Wrote the paper: SS ADW MDB.

## Data availability statement

The data that support the findings of this study will be openly available in the gene expression omnibus (GEO) at [https://www.ncbi.nlm.nih.gov/geo/], reference number [*reference number will be provided upon acceptance of the manuscript*].

