## Supplemental figures 1-3 for "The transcriptional landscape of human microglia reveals strong conservation of miRNAs and preservation of function across vertebrate species"

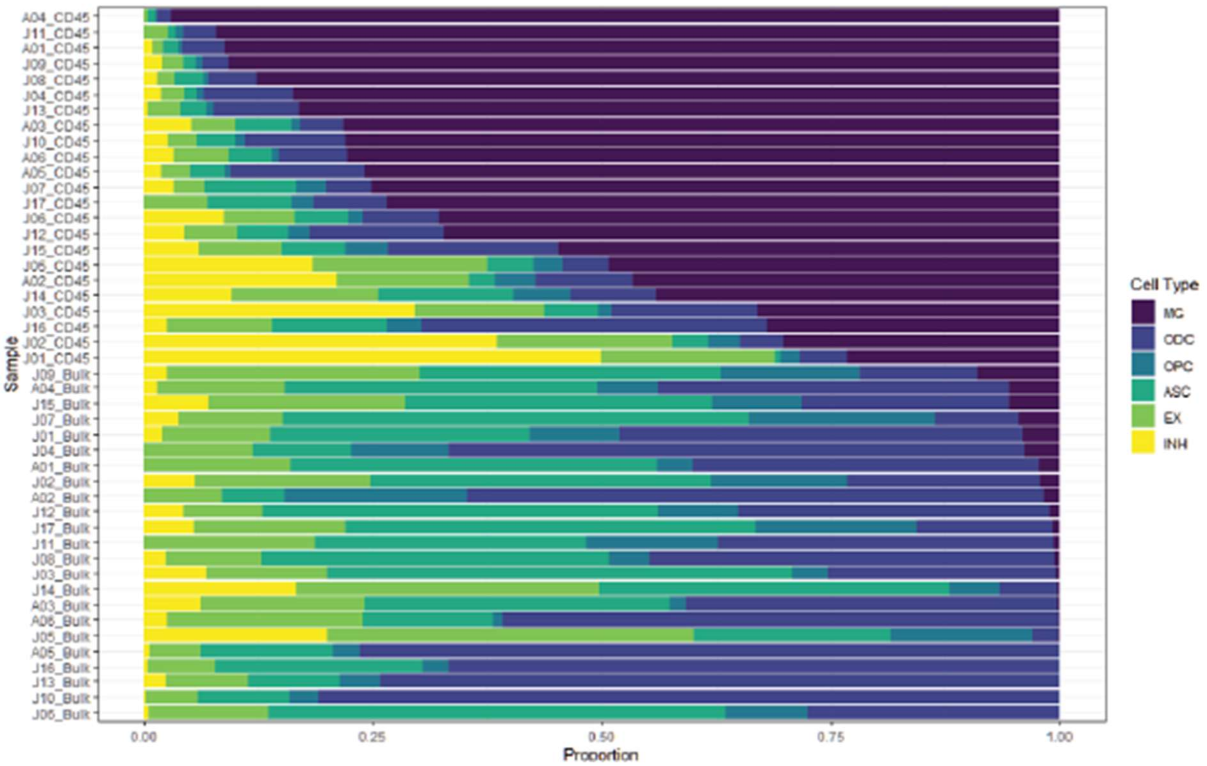


**Supplementary Figure 1. Validation of cell type with CIBERSORTx.** RNA sequencing data from CD45$^{+ve}$ microglial and CD45$^{-ve}$ bulk cell samples were compared against a single nuclear RNAseq dataset derived from human cortical tissue[31] to determine cellular proportions. MG, microglia, ODC, oligodendrocyte, OPC, oligodendrocyte progenitor cell, ASC, astrocyte, Ex, excitatory neurons, INH, inhibitory neurons. N= 23 CD45+ve, 23 CD45-ve.

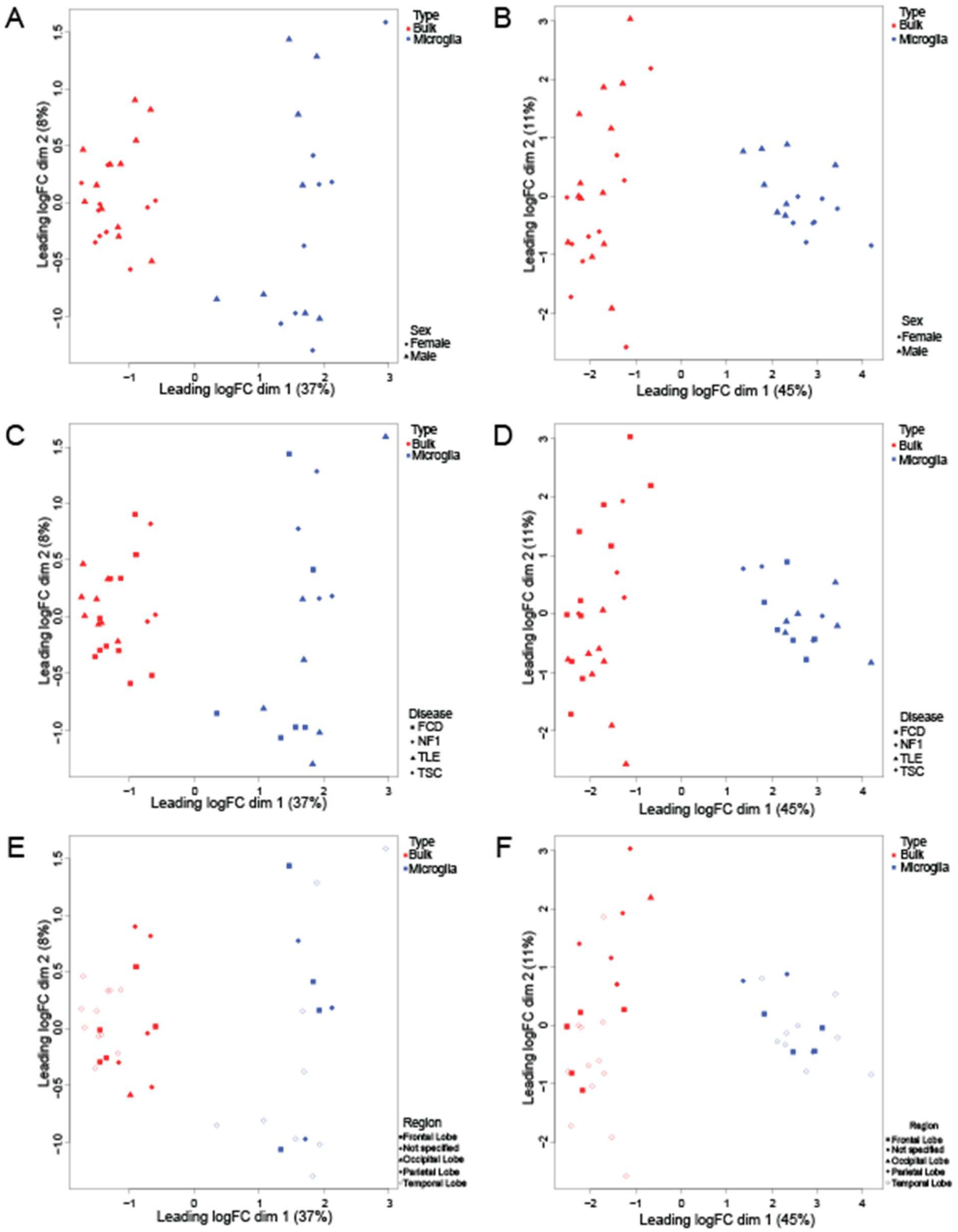


**Supplementary Figure 2. Human miRNA and mRNA expression does not cluster based on sex, disease, or brain region.** MDA plots of analysis of RNA sequencing data with samples labelled by sex (**A**. miRNA, **B**. mRNA), disease (**C**. miRNA, **D**. mRNA), or brain region (**E**. miRNA, **F** mRNA). N= 16 CD45+ve, 22 CD45-ve.

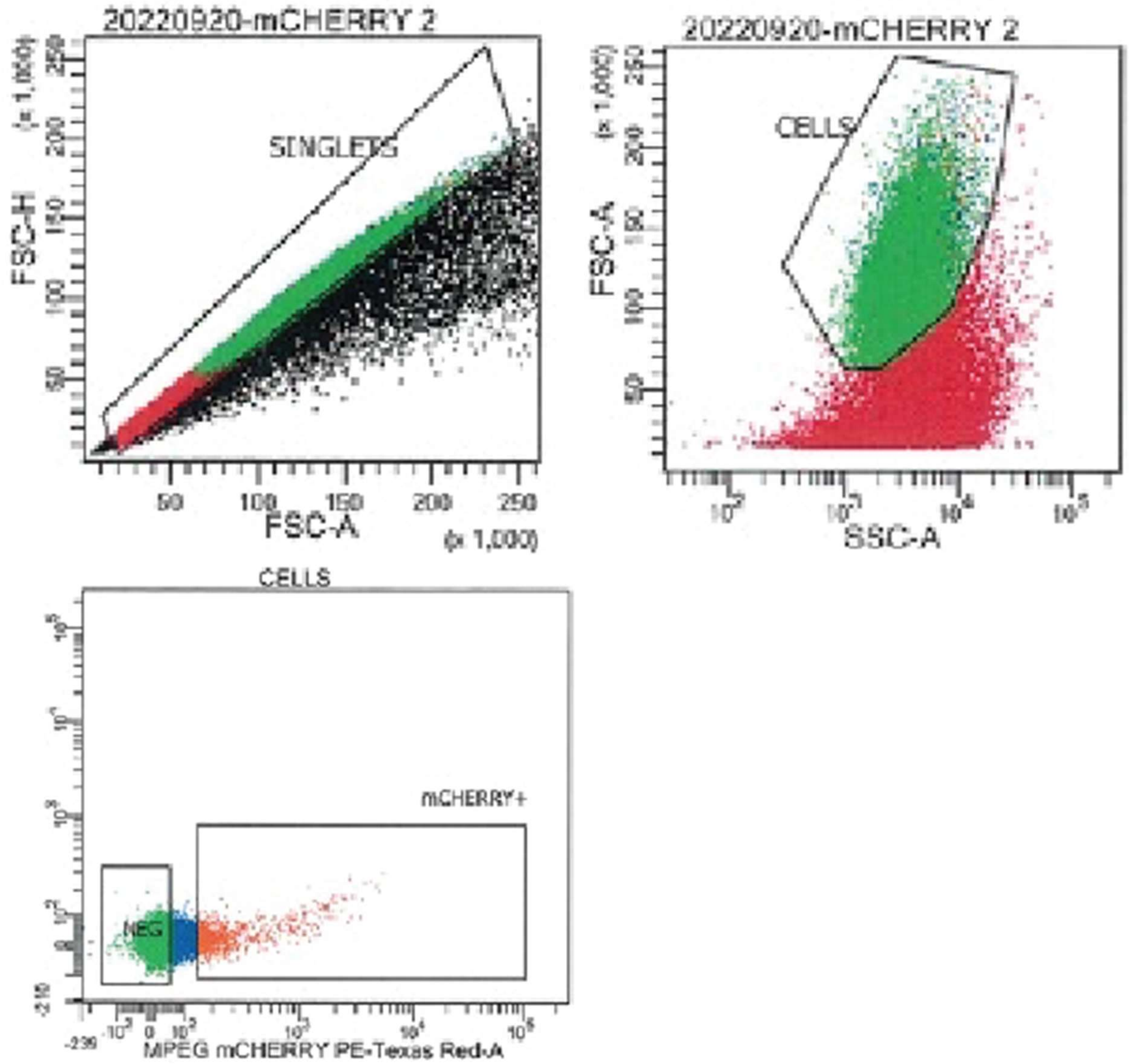


**Supplementary Figure 3. Gating strategy for the purification of mCherry$^{+ve}$ microglia and mCherry$^{-ve}$ bulk brain cells.**
